# Longitudinal associations between task-control network functioning, cognition, and the environment in early adolescence

**DOI:** 10.64898/2026.10.16.759634

**Authors:** K.J. Paige, C. Michael, M.F. Molloy, O. Kardan, M. Angstadt, C. Sripada, L.W. Hyde, M.M. Heitzeg, A.S. Weigard

## Abstract

Neural maturation in the “task-control” frontoparietal and dorsal attention networks is thought to support the development of adolescents’ cognitive and adaptive functioning. Further, theory suggests that family and neighborhood contexts shape brain network development. Yet, few longitudinal studies have examined the co-development of brain network mechanisms with cognitive and adaptive functioning or with environmental factors that could shape the maturation of these mechanisms. The present study addressed these gaps. We used longitudinal data spanning late childhood (*M*age=10 years) to early adolescence (*M*age=12) from the Adolescent Brain Cognitive Development (ABCD) Study (*n*=4862). Stable and construct-valid neural signatures of task-control network functioning were estimated using longitudinal bifactor modeling of neural responses to cognitive demand during the n-back task. Multivariate latent change-score models were used to examine co-development of this signature with cognitive efficiency (drift rate from the diffusion model), teacher-reported attention problems, family conflict, and neighborhood safety. Models accounted for baseline family income. Robust reciprocal relations between task-control activation and cognitive efficiency were identified and the latent change scores for efficiency and task-control activation were positively correlated.

Similar reciprocal associations were observed between cognitive efficiency and attention problems. Baseline family income predicted the rate of change in task-control network functioning. This study demonstrates that task-control network functioning co-develops with cognition, which in turn supports attentional functioning development across early adolescence. Further, findings highlight the utility of this neural signature of task-control network functioning for understanding the neural basis of cognitive development and the mechanisms through which environmental experiences shape brain and behavior.

**Graphical abstract:** 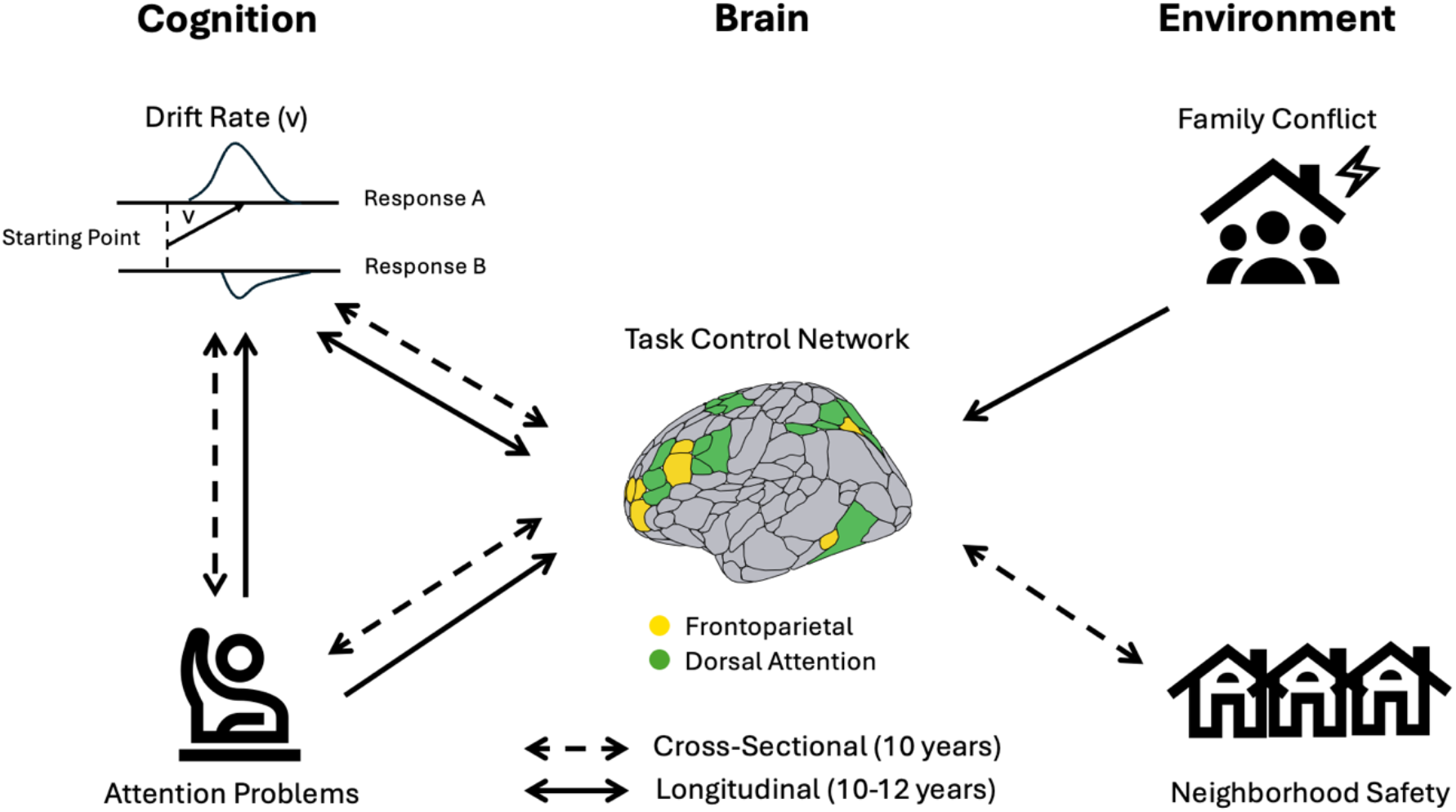

## 1. Introduction

Adolescence is a period of marked development in neurocognitive functions (Blakemore & Choudhury, 2006; Boelema et al., 2014; Larsen & Luna, 2018; Satterthwaite et al., 2013). Importantly, healthy neurocognitive development is critical to concurrent and long-term mental and physical health, as well as future educational attainment and economic stability (Caspi et al., 2013; Hofer & Clouston, 2014; Moffitt et al., 2011; Paige et al., 2021; Strenze, 2007). Thus, a clear understanding of the mechanisms underlying healthy neurocognitive development, as well as environmental factors that may shape it, is critical to public health. The maturation of dorsal and ventral attention and frontoparietal control brain networks have been proposed to play a central role in behavioral maturation of neurocognitive abilities during this developmental period (Dong et al., 2024; Keller et al., 2023). Notably, these brain maturational processes are potentially sensitive to environmental factors (Kardan et al., 2026; Rakesh et al., 2023; Tooley et al., 2021). Characterizing this dynamic brain-behavior co-development and its environmental predictors is of central importance to building a scientific understanding of neurocognitive maturation and for identifying potentially modifiable factors that impact healthy brain development. However, this has been challenging due to a lack of psychometrically stable brain measures within largescale longitudinal neuroimaging studies. The current study addresses this gap.

### 1.1. Neural maturation underlying development of neurocognitive abilities

Longitudinal neuroimaging studies suggest that the frontoparietal and dorsal attention networks are subject to substantial maturation across childhood and early adolescence. Indeed, a recent study investigating the organization of brain networks across ages 6 to 14 years found evidence of progressive maturation in the frontoparietal network, which manifested as reduced modular dynamics (Lei et al., 2022). This is consistent with evidence that developmental trajectories in frontoparietal activity during feedback learning tend to increase from childhood through middle adolescence and peak/level off in late adolescence through early adulthood (Paige et al., 2024; Peters et al., 2016). There is likewise strong evidence for structural and functional changes occurring in the dorsal attention network across childhood and adolescence (Saito et al., 2022; Vinette & Bray, 2015). Importantly, neural functioning across these “task-control” networks is thought to support cognitive performance and behavioral outcomes. Yet, developmental cognitive neuroscience cannot determine how these neural systems develop alongside cognition and behavior unless we can measure their functioning consistently across age. This has been a major challenge: existing measures do not establish whether the same neural construct is being captured consistently across development. We address this gap by utilizing a neural measure of task-control network functioning that is stable across development, shows longitudinal measurement invariance, and demonstrates construct validity (Paige, Molloy, et al., 2026). We apply Latent Change Score modeling (LCS) to these developmentally invariant measures to examine the temporal dynamics between neural, behavioral, and questionnaire measures of cognition **(Figure 1)**.

**Figure 1.**
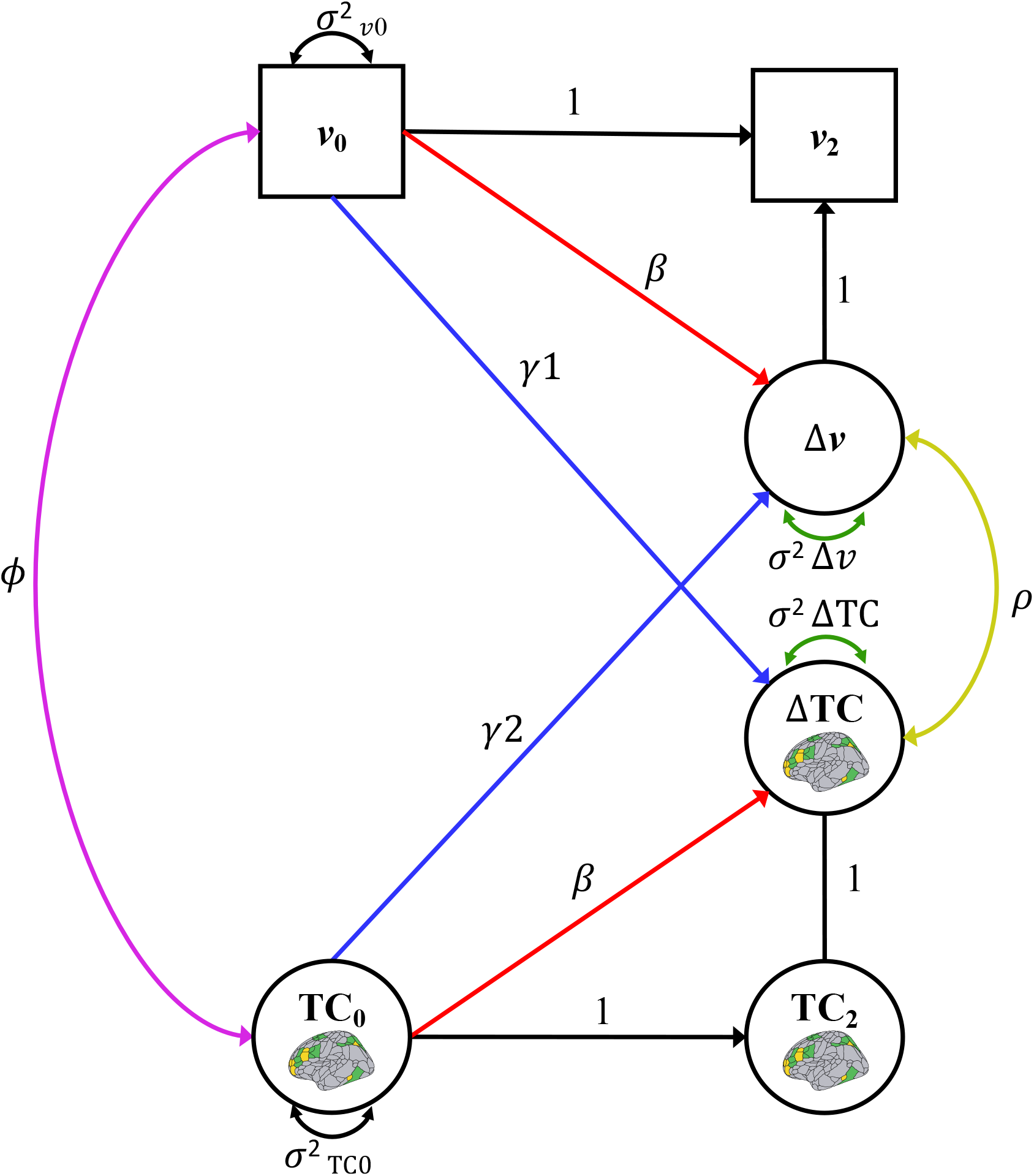
Conceptual figure of a latent change score model examining co-development between general task-control network activation and drift rate. In the current study, this latent change score model was extended to also include teacher-reported attention; we omitted this third variable from the figure for clarity. TC=Task-control network activation. v=cognitive efficiency (drift rate) on n-back task. 0 denotes baseline assessment of ABCD (ages 9-10 years) and 2 denotes year 2 (ages 11-12). Covariate pathways are omitted from figure for clarity but are reported in detail in the text for the univariate model results.

Computational modeling of behavioral task performance has revealed that the flexibility of ‘task-positive’ brain networks’ responses to cognitive demand strongly relates to individual differences in efficiency of evidence accumulation, a basic computational process that supports cognitive performance across diverse tasks (Weigard, Angstadt, et al., 2024). Specifically, individuals with greater efficiency of evidence accumulation tend to have higher activation of the frontoparietal and dorsal attention networks under high cognitive demand but lower activation of the same networks under low demand (Weigard, Angstadt, et al., 2024). This pattern was replicated across the baseline sample of the Adolescent Brain Cognitive Development^SM^ (ABCD) Study of 9- to 10-year-old youth and the Human Connectome Project (HCP) sample of young adults age 22-35 years. A recent study built on these findings, developing a psychometrically sound neural signature of task-control network functioning across HCP and the first two neuroimaging assessments of ABCD (Paige, Molloy, et al., 2026). Findings revealed moderate to strong within-time correlations between higher neural activation across the task-control networks (spanning the frontoparietal and dorsal attention networks) and better behavioral performance across a wide variety of cognitive tasks. Higher activation on this general task-control activation factor was also concurrently related to fewer attention problems in late childhood and early adolescence. Critically, the task-control network factor showed evidence of strict measurement invariance across ABCD timepoints, allowing for quantitative comparisons of individuals’ levels of network activation across ages.

These studies, however, focused on cross-sectional associations rather than the more complex task of characterizing the co-development of task-control network functioning with cognitive and adaptive function in longitudinal data. Although characterizing the co-development of neural-level metrics with behavior in largescale samples like ABCD is critical for understanding which brain features support cognitive and behavioral development, we are unaware of any prior work that has applied developmental modeling methods, such as LCS models, to neuroimaging measures of neurocognition in ABCD. However, one prior longitudinal study demonstrates the co-development of cognitive efficiency (estimated from a different ABCD task, the stop-signal) and attention problems across early adolescence (Wiker et al., 2024). Taken together with cross-sectional associations from neuroimaging studies, these findings suggest that maturation in task-control networks may support improvements in cognitive efficiency and attentional functioning seen across this developmental period. However, due to a lack of longitudinal neuroimaging studies and age-invariant measures of task-control network functioning, the temporal dynamics of how this brain feature may co-develop with behavioral and adaptive functioning remain poorly understood.

### 1.2. Environmental experiences and neurocognitive development

Decades of cross-species research have established that environmental adversity is associated with individual differences in neurocognitive development. Adversity exposure can occur across levels of context, from proximal household to more distal neighborhood and school contexts (Bronfenbrenner, 2000), and can influence neurodevelopment through multiple mechanisms, such as stress and activity-dependent synaptic remodeling (Sheridan & McLaughlin, 2022). A broad literature has documented that family conflict and neighborhood threat are important forms of environmental adversity that may shape neurocognitive development (Andrews et al., 2021; Lanjekar et al., 2022; Lewis et al., 2025; McLaughlin et al., 2019). Indeed, perceived social threats across family and neighborhood contexts have been associated with altered functional connectivity patterns across frontoparietal and dorsal attention networks (Tsomokos et al., 2025; Wen et al., 2023). Importantly, recent neurodevelopmental frameworks propose that higher-order association cortex implicated in cognitive control may express peak plasticity during adolescence, suggesting that task-control networks may be particularly susceptible to adversity during this developmental window (Larsen et al., 2023; Michael et al., 2025; Sydnor et al., 2021). Together, these findings indicate that environmental adversity across family and neighborhood contexts may influence key brain systems underlying the development of goal-directed cognition. Yet, here again, past work is limited by a lack of neural measures that demonstrate invariance and stability across time. Without such a measure, it remains difficult to determine whether observed associations reflect the impact of the environment on adolescent brain development or baseline associations related to factors that precede adolescence.

Previous cross-sectional research has commonly interpreted associations between adversity exposure and neural phenotypes as reflecting potential environmental influences on brain maturation. Although this causal pathway is supported by experimental non-human models which demonstrate altered neurochemical signaling in prefrontal task control regions following chronic stress exposure (Perica & Luna, 2023), alternative interpretations are plausible. Seminal developmental theories, such as transactional and developmental psychopathology frameworks, emphasize dynamic and reciprocal interactions among developmental pathways, environmental experiences, and child characteristics (Sameroff, 2009; Sroufe & Rutter, 1984). Thus, whereas adversity exposure may influence neurocognitive development, neurobehavioral characteristics of the child may also engender changes in their environment (Hyde et al., 2024). Within this framework, prior longitudinal work has shown that variability in cognition and self-regulation is associated with changes in parenting across childhood and adolescence (Blair et al., 2014; Khurana et al., 2024). Even though individual differences in cognition partly stem from variability in underlying neural substrates, to date, these developmental transactions are almost exclusively supported by behavioral evidence. Put differently, limited longitudinal research has characterized reciprocal interactions among family and neighborhood contexts and neuroimaging indicators of neurocognitive maturation during adolescence, a period characterized by rapid neurocognitive remodeling in the context of dynamically changing environmental demands.

### 1.3. The current study

Adolescence is a period of substantial maturation in task-control networks that are thought to support cognitive and attentional functions. Yet, there is no longitudinal research characterizing the ways in which task-control network activation co-develops with behavioral measures of cognition and attention problems. Such research requires psychometrically sound and age-invariant measures of task-control network functioning. Further, leading developmental cognitive neuroscience theories suggest that environmental experiences shape neural development across adolescence, but a lack of psychometrically sound longitudinal neuroimaging measures has presented a significant challenge for empirical studies. The current study addresses these gaps. We use longitudinal neuroimaging, behavioral, and questionnaire data from late childhood (9–10 years) to early-adolescence (11–13 years) from ABCD. Neural signatures of general task-control network functioning were computed and diffusion modelling was applied to behavioral data from the n-back task to measure efficiency of evidence accumulation. This approach allows us, for the first time, to 1) examine how task-control network maturation co-develops with cognitive performance and attention problems and, 2) investigate how family and neighborhood contexts may shape task-control network maturation across a critical period of brain development, early adolescence.

#### 1.3.1. Hypotheses

We hypothesize that neural maturation in task-control networks will co-develop with maturation in behavioral measures of cognitive efficiency and questionnaire measures of attentional functioning. Given prior work demonstrating the co-development of cognitive efficiency and attention problems (Wiker et al., 2024), we also predict that cognitive efficiency will show evidence of co-development with attention problems, such that maturation in cognitive efficiency will be related to decreases in attention problems. Further, we expect that low family conflict and high perceived neighborhood safety will associate with improved task-control network maturation.

## 2. Method

### 2.1. Participants

Data were drawn from ABCD Release 5.1 (http://dx.doi.org/10.15154/z563-zd24). Here, we included data from baseline (*M*age=10.01 years; *n*=4862) and Year 2 (*M*age=11.99; *n*=3648). These sample sizes were largely determined by the subset of the sample who completed the neuroimaging protocol and had usable neuroimaging data (see inclusion criteria below).

### 2.2. Procedure

The ABCD Study® is a large, multisite study with multimodal data obtained from over 11,000 adolescents collected from 21 sites across the United States (Casey et al., 2018; Garavan et al., 2018). The study conforms to the rules and procedures of each site’s Institutional Review Board (IRB). The University of California San Diego’s IRB serves as a single IRB for the ABCD consortium (IRB#: 160091), and the IRBs of each of the 21 sites also approve the study. All participants provide informed consent (parents) or informed assent (children). High resolution structural images (1.0mm3 isotropic voxels) T1-weighted and T2-weighted), diffusion weighted imaging, and multiple functional MRI runs (TR=800 ms, TE=30 ms, 2.4 mm isotropic voxels) were collected for all participants.

### 2.3. Measures

#### 2.3.1. N-back task

Task-control network activation was measured using the ABCD emotional N-back (EN-Back) task during high cognitive load (2-back versus 0-back contrast). Participants were shown a series of stimuli that included places and happy, fearful, and neutral faces in 10-trial blocks and were asked to response as to whether each stimulus was a target in each of two conditions. In the 0-back conditions targets matched images that were displayed at the beginning of the block. In the 2-back, targets were defined as stimuli that matched the stimulus presented two spaces back in the sequence. fMRI data with 2.4 mm isotropic voxels (TR=800ms, TE=30ms) were collected concurrently during task performance. The full task comprised 2 imaging runs of 8 blocks (four 2-back, four 0-back) each. Additional details about the task design and acquisition can be found in Casey et al., 2018.

#### 2.3.2. Teacher-reported attention problems

Attention problems were assessed by six items on the Brief Problem Monitor (Achenbach, T.M., 2009), which was administered to teachers at baseline and year 2. Caregivers were asked to choose the teacher who had the most frequent contact with their child (Barch et al., 2021). Example items include “can’t concentrate or pay attention for long” and, “can’t sit still, restless or hyperactive.” Raw scores of attention problems were used in the current study. Cronbach’s α values were good (baseline α=0.89; year 2 α=0.89).

#### 2.3.3. Family conflict

The Family Conflict Scale, adapted from the Family Environment Scale (Moos, R.H., & Moos, B.S., 1994; Sanford et al., 1999), was administered to both youth and their caregivers. This questionnaire contains 9 items assessing openly expressed conflict among family members. Example items included, “We fight a lot in our family,” and “Family members sometimes get so angry they throw things.” Responses were binary (0=“No”, 1=“Yes”). We used sum scores of family conflict for both youth and caregiver report. Cronbach’s α values were acceptable for youth (baseline α=0.68; year 2 α=0.64) and caregiver (baseline α=0.66; year 2 α=0.68) reports.

#### 2.3.4. Perceived neighborhood safety

Participants completed the Neighborhood Safety and Crime questionnaire at baseline and year 2, a one question, youth-report measure asking the participant to rate their agreement with the statement “My neighborhood is safe from crime” on a Likert rated scale. Responses were coded such that 1=strongly disagree and 5=strongly agree (Mujahid et al., 2007; PhenX Toolkit, n.d.).

#### 2.3.5. Covariates

Covariates included mean framewise displacement (FD) at baseline and year 2, baseline age, and baseline family income. Family income was categorized as low (<$50,000), middle ($50,000–$99,999), or high (≥$100,000). Low-and high-income groups were dummy coded relative to the middle-income group, which served as the reference category for all analyses.

### 2.4. Neuroimaging preprocessing and ROI extraction

Neuroimaging exclusion criteria, data quality checks, preprocessing, and analyses are described in detail in (Weigard, Angstadt, et al., 2024). Briefly, preprocessing was performed using fMRIPrep v1.5.0 (Esteban et al., 2019). Structural T1-weighted (T1w) and T2-weighted images were run through recon-all using FreeSurfer v6.0.1. Functional images were then corrected for fieldmap distortions, rigidly coregistered to the T1, motion corrected, normalized to standard space, and transformed to CIFTI space.

Task data were analyzed following HCP scripts for both datasets. Models were constructed using FSL, averaging across runs, smoothed (2mm FWHM Gaussian), and highpass filtered. First-level models included nuisance parameters for motion, white matter, cerebrospinal fluid, and framewise displacement. Each condition (2-back vs. 0-back) and category of stimuli (e.g., places) were modeled separately, then averaged across stimuli for the final relevant 2back-0back contrast. Contrast values were then averaged within each region in the 100-parcel Schaeffer parcellation to reduce the dimensionality for factor analysis. All frontoparietal and dorsal attention regions from this parcellation were retained for analysis and activation data were averaged withing each parcel. Several parcels in in midline areas of the brain were averaged across hemispheres because they exhibited strong correlations with one another. A total of 21 variables were included across the task-control network, 11 from the frontoparietal network and 10 from the dorsal attention network. Bifactor models of task-control network activation were specified at each assessment, with all indicators loading onto a general factor, 11 indicators additionally loading onto an frontoparietal-specific factor, and 10 additionally loading onto a dorsal attention-specific factor. This factor structure demonstrated strict longitudinal measurement invariance across development. Additionally, the general factor of task control-network activation demonstrated robust construct validity. All details are provided in (Paige, Molloy, et al., 2026). Thus, in the current study, we utilize the general factors of task-control network activation to investigate its co-development with maturation in behavioral measures of cognition and questionnaire measures of attentional functioning, as well as measures of adversity across family and neighborhood contexts.

### 2.5. Data analytic strategy

#### 2.5.1. Diffusion decision model

The diffusion decision model (DDM) (Ratcliff, 1978; Ratcliff et al., 2016) was applied to the n-back task to obtain estimates of individuals’ efficiency of evidence accumulation (EEA). We used a recent version of the DDM designed to parametrically separate EEA for goal-relevant information from individuals’ strategic use of goal-irrelevant familiarity evidence to complete the task, which is important for describing individual differences in performance on the ABCD task (Weigard, Suzuki, et al., 2024). Parameter estimation procedures were identical to those that were implemented, and described in detail, in this prior study (Weigard, Suzuki, et al., 2024). Briefly, all models were estimated in the Dynamic Models of Choice R suite (Heathcote et al., 2019) using individual-level Bayesian estimation via differential-evolution Markov chain Monte Carlo sampling (Turner et al., 2013). Prior to estimation, trials with response times <200ms were excluded as fast guesses and individuals with accuracy rates <0.55 or omission (i.e., non-response) rates >0.25 on either the 0-or 2-back task were excluded due to likely disengagement from the task. Hierarchical Bayesian versions of the n-back model were first fit to a randomly selected subsample of 200 singleton ABCD participants to generate informative priors for the baseline and year 2 waves, and these individuals were then excluded from further analyses involving DDM parameters. Individual-level Bayesian estimation using the resulting informative priors was then conducted for the rest of the ABCD sample separately at baseline and year 2 waves. Convergence was determined by the Gelman-Rubin statistic falling below 1.10 (Gelman & Rubin, 1992) and model fit was assessed using posterior predictive plots. Medians of individuals’ posterior distributions of the EEA parameter were then retained as point estimates for analyses. All procedures, initial prior distributions, and posterior predictive plots are detailed in the open access prior article and its supplement (Weigard, Suzuki, et al., 2024).

#### 2.5.2. Latent change score models

Data preparation and descriptive analyses were conducted in R/R Studio (v4.2.3; R Core Team 2023; Hallquist & Wiley, 2018).

LCS (also referred to as latent difference scores) models within a structural equation modeling framework were used to examine the temporal dynamics between neural, behavioral, and questionnaire measures of cognition and environmental adversity (e.g., **Figure 1**). The LCS *Δi* captures the difference for individual *i* between Times 1 and 2. LCS models decompose continuous variables into two components, the continuous latent “true” score and a residual measurement error term. As outlined by de Haan and colleagues (2018), the first step when estimating an LCS model with separate constructs is to assess for measurement invariance. Testing longitudinal measurement invariance assesses whether the difference score has the same meaning across time (McArdle, 2009). Our neural signature of task-control network functioning demonstrated strict invariance across baseline and year 2 in ABCD (Paige, Molloy, et al., 2026). We did not examine longitudinal measurement invariance for observed variables. Next, we estimated univariate LCS models to assess mean change in each construct. Then, we tested multivariate LCS models to examine co-development of brain, behavioral, and questionnaire metrics of cognition, as well as co-development of brain and environment variables. In other words, LCS models provide a framework for characterizing developmental change across two repeated measures and testing predictors of that change. Head motion, baseline age, and baseline family income were included as covariates. Correlations among covariates were allowed. A conceptual figure of a multivariate LCS model is depicted in **Figure 1**.

All structural equation models were specified in Mplus 8.8 using Maximum likelihood with robust standard errors (MLR) and a Full-information Maximum Likelihood estimator (FIML) to account for missing data (Muthén, L.K., & Muthén, B.O., 1998). Youths were clustered within families and stratified by study site using the CLUSTER IS and STRATIFICATION IS statements, respectively. Model fit was assessed chi-square (smaller values indicate better fit), Comparative Fit Index (CFI), Root Mean Square Error of Approximation (RMSEA), and Standardized Root Mean Square Residual (SRMR). CFI>0.90, RMSEA≤0.05, and SRMR<0.06 indicate acceptable model fit (Hu & Bentler, 1999).

## 3. Results

### 3.1. Univariate latent change score models

#### 3.1.1. Task-control network activation and drift rate increase and attention problems decrease across early adolescence

The univariate LCS models all provided an acceptable fit to the data **(*see* Table 1)**. Results from univariate LCS models are reported in **Table 2**. The models showed significant positive mean change in task-control network functioning and drift rate, indicating that task-control network activation and cognitive efficiency demonstrate quantitative increases across ages 10 to 12. The univariate LCS model for teacher-reported attention problems suggested that attention problems decreased across ages 10 to 12, on average. Univariate LCS models indicated decreases in caregiver-and youth-reported family conflict, and increases in perceived neighborhood safety across ages 10 to 12. There were also significant variances in baseline scores and change for all variables of interest. The significant negative correlations between baseline and change indicated that individuals with high initial levels of each variable (e.g., task-control network activation) had slower increases in that same variable across ages 10 to 12.

**Table 1.**
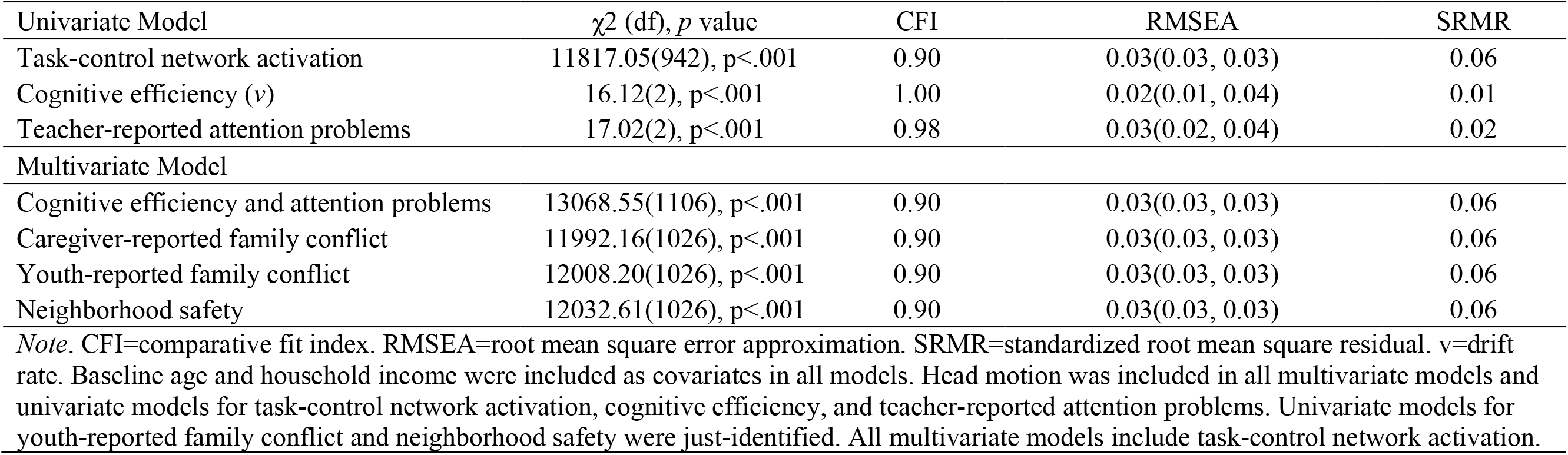
Fit statistics for latent change score models.

| Univariate Model | $\chi^2$ (df), *p* value | CFI | RMSEA | SRMR |
| --- | --- | --- | --- | --- |
| Task-control network activation | 11817.05(942), *p*<.001 | 0.90 | 0.03(0.03, 0.03) | 0.06 |
| Cognitive efficiency (*v*) | 16.12(2), *p*<.001 | 1.00 | 0.02(0.01, 0.04) | 0.01 |
| Teacher-reported attention problems | 17.02(2), *p*<.001 | 0.98 | 0.03(0.02, 0.04) | 0.02 |
| Multivariate Model |  |  |  |  |
| Cognitive efficiency and attention problems | 13068.55(1106), *p*<.001 | 0.90 | 0.03(0.03, 0.03) | 0.06 |
| Caregiver-reported family conflict | 11992.16(1026), *p*<.001 | 0.90 | 0.03(0.03, 0.03) | 0.06 |
| Youth-reported family conflict | 12008.20(1026), *p*<.001 | 0.90 | 0.03(0.03, 0.03) | 0.06 |
| Neighborhood safety | 12032.61(1026), *p*<.001 | 0.90 | 0.03(0.03, 0.03) | 0.06 |

*Note.* CFI=comparative fit index. RMSEA=root mean square error approximation. SRMR=standardized root mean square residual. *v*=drift rate. Baseline age and household income were included as covariates in all models. Head motion was included in all multivariate models and univariate models for task-control network activation, cognitive efficiency, and teacher-reported attention problems. Univariate models for youth-reported family conflict and neighborhood safety were just-identified. All multivariate models include task-control network activation.

**Table 2.** Results from univariate latent change score models.

| Parameter | Model |  |  |  |  |  |  |  |  |  |  |  |
| --- | --- | --- | --- | --- | --- | --- | --- | --- | --- | --- | --- | --- |
|  | TC network activation |  | Cognitive efficiency (*v*) |  | Teacher-rep. attention prob. |  | Caregiver-rep. family conflict |  | Youth-rep. family conflict |  | Neighborhood safety |  |
|  | Est. | SE | Est. | SE | Est. | SE | Est. | SE | Est. | SE | Est. | SE |
| Baseline mean | 20.47 | 0.43 | 0.91 | 0.004 | 2.84 | 0.05 | 2.54 | 0.02 | 2.05 | 0.02 | 4.03 | 0.01 |
| Mean change | 15.78 | 0.77 | 0.55 | 0.01 | -1.14 | 0.16 | -0.10 | 0.04 | -0.12 | 0.02 | 0.06 | 0.02 |
| Variance of baseline score | 342.18 | 13.25 | 0.15 | 0.002 | 10.42 | 0.24 | 3.83 | 0.06 | 3.82 | 0.05 | 1.20 | 0.02 |
| Variance of change score | 401.92 | 22.21 | 0.13 | 0.003 | 7.85 | 0.41 | 3.34 | 0.06 | 4.45 | 0.07 | 1.57 | 0.03 |
| Correlation of baseline-change | -0.51 | 0.02 | -0.34 | 0.01 | -0.53 | 0.02 | -0.46 | 0.01 | -0.60 | 0.01 | -0.67 | 0.01 |
| Baseline age predicting change | 0.08 | 0.02 | 0.08 | 0.01 | 0.03 | 0.02 | 0.001 | 0.002 | 0.04 | 0.008 | -0.02 | 0.007 |
| Low income predicting change | 0.01 | 0.02 | -0.10 | 0.02 | 0.07 | 0.02 | -0.02 | 0.05 | 0.06 | 0.01 | -0.10 | 0.01 |
| High income predicting change | 0.06 | 0.02 | 0.08 | 0.02 | -0.05 | 0.02 | 0.08 | 0.04 | -0.03 | 0.01 | 0.07 | 0.009 |

*Note.* TC=Task-control. Rep.=reported. Prob.=Problems. *v*=drift rate.

#### 3.1.2. Baseline household income predicts change in task-control network activation, drift rate, and attention problems

Zero-order correlations for observed variables are provided in **Table 3**. Univariate LCS models revealed associations between baseline covariates and longitudinal change in study variables **(Table 2)**. Specifically, older baseline age predicted greater growth in task-control network activation and drift rate. Relative to youth from households earning $50,000–$99,999, youth from households earning less than $50,000 displayed smaller increases in drift rate and smaller decreases in attention problems. Conversely, youth from households earning $100,000 or more showed larger increases in task-control network activation and drift rate, and faster decreases in attention problems. Regarding family conflict and perceived neighborhood safety, older baseline age predicted smaller decreases in youth-reported family conflict and slower increases in perceived neighborhood safety. Relative to youth from households earning $50,000–$99,999, youth from households earning less than $50,000 displayed smaller decreases in youth-reported family conflict and smaller increases in perceived neighborhood safety. Youth from households earning $100,000 or more showed larger decreases in youth-reported family conflict and greater growth in perceived neighborhood safety.

**Table 3.**
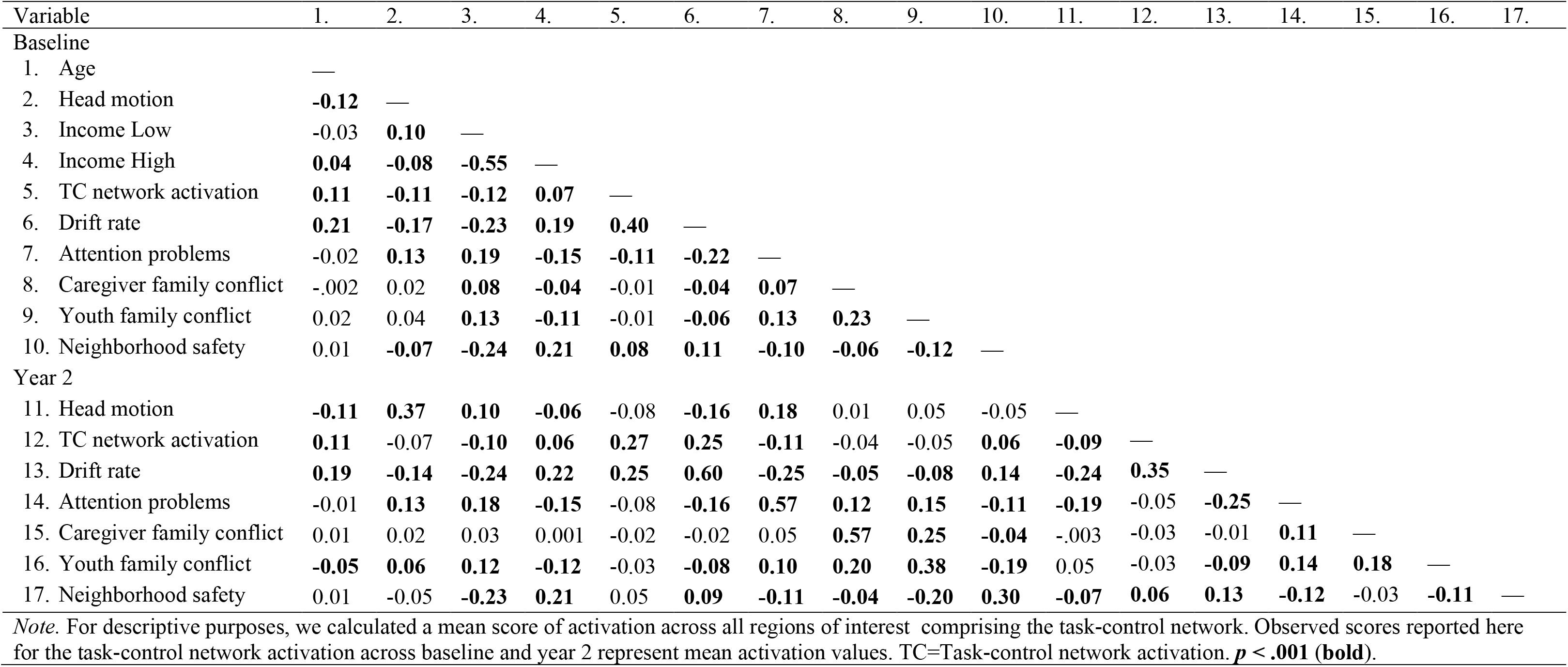
Bivariate correlations for observed variables and covariates.

| Variable | 1. | 2. | 3. | 4. | 5. | 6. | 7. | 8. | 9. | 10. | 11. | 12. | 13. | 14. | 15. | 16. | 17. |
| --- | --- | --- | --- | --- | --- | --- | --- | --- | --- | --- | --- | --- | --- | --- | --- | --- | --- |
| Baseline |  |  |  |  |  |  |  |  |  |  |  |  |  |  |  |  |  |
| 1. Age | — |  |  |  |  |  |  |  |  |  |  |  |  |  |  |  |  |
| 2. Head motion | **-0.12** | — |  |  |  |  |  |  |  |  |  |  |  |  |  |  |  |
| 3. Income Low | -0.03 | **0.10** | — |  |  |  |  |  |  |  |  |  |  |  |  |  |  |
| 4. Income High | **0.04** | **-0.08** | **-0.55** | — |  |  |  |  |  |  |  |  |  |  |  |  |  |
| 5. TC network activation | **0.11** | **-0.11** | **-0.12** | **0.07** | — |  |  |  |  |  |  |  |  |  |  |  |  |
| 6. Drift rate | **0.21** | **-0.17** | **-0.23** | **0.19** | **0.40** | — |  |  |  |  |  |  |  |  |  |  |  |
| 7. Attention problems | -0.02 | **0.13** | **0.19** | **-0.15** | **-0.11** | **-0.22** | — |  |  |  |  |  |  |  |  |  |  |
| 8. Caregiver family conflict | -.002 | 0.02 | **0.08** | **-0.04** | -0.01 | **-0.04** | **0.07** | — |  |  |  |  |  |  |  |  |  |
| 9. Youth family conflict | 0.02 | 0.04 | **0.13** | **-0.11** | -0.01 | **-0.06** | **0.13** | **0.23** | — |  |  |  |  |  |  |  |  |
| 10. Neighborhood safety | 0.01 | **-0.07** | **-0.24** | **0.21** | **0.08** | **0.11** | **-0.10** | **-0.06** | **-0.12** | — |  |  |  |  |  |  |  |
| Year 2 |  |  |  |  |  |  |  |  |  |  |  |  |  |  |  |  |  |
| 11. Head motion | **-0.11** | **0.37** | **0.10** | **-0.06** | -0.08 | **-0.16** | **0.18** | 0.01 | 0.05 | -0.05 | — |  |  |  |  |  |  |
| 12. TC network activation | **0.11** | -0.07 | **-0.10** | **0.06** | **0.27** | **0.25** | **-0.11** | -0.04 | -0.05 | **0.06** | **-0.09** | — |  |  |  |  |  |
| 13. Drift rate | **0.19** | **-0.14** | **-0.24** | **0.22** | **0.25** | **0.60** | **-0.25** | **-0.05** | **-0.08** | **0.14** | **-0.24** | **0.35** | — |  |  |  |  |
| 14. Attention problems | -0.01 | **0.13** | **0.18** | **-0.15** | -0.08 | **-0.16** | **0.57** | **0.12** | **0.15** | **-0.11** | **-0.19** | -0.05 | **-0.25** | — |  |  |  |
| 15. Caregiver family conflict | 0.01 | 0.02 | 0.03 | 0.001 | -0.02 | -0.02 | 0.05 | **0.57** | **0.25** | **-0.04** | -.003 | -0.03 | -0.01 | **0.11** | — |  |  |
| 16. Youth family conflict | **-0.05** | **0.06** | **0.12** | **-0.12** | -0.03 | **-0.08** | **0.10** | **0.20** | **0.38** | **-0.19** | 0.05 | -0.03 | **-0.09** | **0.14** | **0.18** | — |  |
| 17. Neighborhood safety | 0.01 | -0.05 | **-0.23** | **0.21** | 0.05 | **0.09** | **-0.11** | **-0.04** | **-0.20** | **0.30** | **-0.07** | **0.06** | **0.13** | **-0.12** | -0.03 | **-0.11** | — |

Note. For descriptive purposes, we calculated a mean score of activation across all regions of interest comprising the task-control network. Observed scores reported here for the task-control network activation across baseline and year 2 represent mean activation values. TC=Task-control network activation. $p < .001$ (bold).

### 3.2. Multivariate latent change score models

All bivariate models examining associations with task-control network activation provided an acceptable fit to the data **(Table 1)**. Results from all multivariate LCS models are presented in **Tables 4 and5**. With respect to associations with covariates, associations in multivariate models were consistent with those reported above from univariate models.

**Table 4.** Results from multivariate latent change score model examining associations between change in task-control network functioning, cognitive efficiency, and attention problems.

| Parameter | $\beta$ | $SE$ | $p$ |
| --- | --- | --- | --- |
| TC0 ↔ v0 ($\phi_1$) | 0.53 | 0.01 | <.001 |
| TC0 ↔ att0 ($\phi_2$) | -0.12 | 0.02 | <.001 |
| v0 ↔ att0 ($\phi_3$) | -0.18 | 0.02 | <.001 |
| v0 → ΔTC ($\gamma_1$) | 0.09 | 0.03 | 0.001 |
| TC0 → Δv ($\gamma_2$) | 0.08 | 0.02 | 0.002 |
| att0 → ΔTC ($\gamma_3$) | -0.07 | 0.03 | 0.013 |
| TC0 → Δatt ($\gamma_4$) | -0.05 | 0.04 | 0.184 |
| att0 → Δv ($\gamma_5$) | -0.13 | 0.02 | <.001 |
| v0 → Δatt ($\gamma_6$) | -0.02 | 0.03 | 0.555 |
| ΔTC ↔ Δv ($\rho_1$) | 0.32 | 0.02 | <.001 |
| ΔTC ↔ Δatt ($\rho_2$) | 0.01 | 0.04 | 0.817 |
| Δv ↔ Δatt ($\rho_3$) | -0.08 | 0.03 | 0.002 |

Note. The parameters of interest are provided: ( $\phi$ ) correlation at baseline, ( $\gamma$ ) regression and, ( $\rho$ ) correlation of change scores. 0 denotes baseline assessment. TC=Task-control. v=Drift rate (cognitive efficiency). att=Attention problems. Covariate pathways are omitted from tables for clarity but are reported in detail in the text for the univariate model results.

**Table 5.** Results from bivariate latent change score models.

| Parameter | Model |  |  |  |  |  |  |  |  |
| --- | --- | --- | --- | --- | --- | --- | --- | --- | --- |
|  | Caregiver-reported family conflict |  |  | Youth-reported family conflict |  |  | Neighborhood safety |  |  |
| | $\beta$ | $SE$ | $p$ | $\beta$ | $SE$ | $p$ | $\beta$ | $SE$ | $p$ |
| TC0 ↔ z0 ($\phi$) | -0.01 | 0.02 | 0.512 | -0.02 | 0.02 | 0.234 | 0.06 | 0.02 | 0.001 |
| z0 → ΔTC ($\gamma_1$) | -0.03 | 0.02 | 0.095 | -0.01 | 0.02 | 0.534 | 0.02 | 0.02 | 0.291 |
| TC0 → Δz ($\gamma_2$) | 0.00 | 0.02 | 0.997 | -0.01 | 0.01 | 0.571 | 0.02 | 0.01 | 0.077 |
| ΔTC ↔ Δz ($\rho$) | -0.02 | 0.02 | 0.445 | -0.02 | 0.02 | 0.409 | 0.01 | 0.02 | 0.694 |

*Note.* z represents the variable accompanying task-control network activation in the bivariate latent change score model. The four parameters of interest are provided: ( $\phi$ ) correlation at baseline, ( $\gamma_1$ ) regression (baseline z variable predicting change in task-control network functioning), ( $\gamma_2$ ) regression (baseline task-control network functioning predicting change in z variable) and, ( $\rho$ ) correlation of change scores. 0 denotes baseline assessment. TC=Task-control. Results depicted are from models including head motion, baseline age, and baseline family income as covariates. Covariate pathways are omitted from tables for clarity but are reported in detail in the text for the univariate model results.

#### 3.2.1. Task-control network activation, cognitive efficiency (*v*; drift rate), and teacher-reported attention problems co-develop across early adolescence

At baseline, higher task-control network activation was related to faster drift rate with a large effect size. Further, higher task-control network activation at baseline predicted improvement in drift rate across ages 10 to 12, and vice versa; faster baseline drift rate predicted increases in task-control network activation across time. The change scores were also significantly associated, such that improvement in drift rate was related to increases in task-control network activation across ages 10 to 12 with the effect size in the range typically considered “moderate”.

At baseline, higher task-control network activation was related to fewer attention problems. Further, fewer attention problems at baseline predicted increases in task-control network activation across ages 10 to 12. The reciprocal association between baseline task-control network activation and change in attention problems was in the expected direction, but not significant. The change scores were not significantly associated.

Finally, there were several statistically significant associations between drift rate and attention problems. At baseline, faster drift rate was related to fewer attention problems. Fewer attention problems at baseline predicted improvements in drift rate across ages 10 to 12. The reciprocal association between baseline drift rate and change in attention problems was in the expected direction, but not significant. The change scores were significantly associated, such that improvement in drift rate was related to decreases in attention problems across ages 10 to 12.

Associations between task-control network functioning, drift rate, and parent-reported attention problems were consistent with those reported here and are provided in **Supplementary Table 1**.

#### 3.2.2. Family conflict is not associated with task-control network activation

No associations between task-control network activation and family conflict were statistically significant across caregiver and youth-reports. The direction of all associations was negative.

#### 3.2.3. Perceived neighborhood safety is associated with high task-control network activation at age 10

At baseline, greater perceived neighborhood safety was significantly associated with greater task-control network activation, above and beyond associations family income. This indicates that, on average, when 10-year-olds perceived high levels of neighborhood safety they also demonstrated high task-control network activation. No associations with change scores for either neighborhood safety or task-control network activation were statistically significant.

## 4. Discussion

This study was the first to examine the co-development of task-control network functioning with behavioral and questionnaire metrics of neurocognitive development using psychometrically sound, age-invariant neuroimaging measures and a large, population-based longitudinal sample spanning late childhood to early adolescence. Further, we examined how family and neighborhood contexts may shape the neural underpinnings of cognition across a critical period of brain development. Our main findings showed that task-control network functioning co-develops with increases in cognitive efficiency and decreases in attention problems across early adolescence. Further, high perceived neighborhood safety was associated with more task-control network activation at baseline and high household income predicted greater maturation of task-control network activation across early adolescence.

### 4.1. Co-development of brain, behavioral, and clinical measures of cognition

Task-control network activation and behaviorally measured cognitive efficiency increased with age, and teacher-reported attention problems decreased across time. We also found variance in both baseline scores and change scores for all three measures of neurocognition. These findings are consistent with a robust literature documenting reliable individual differences and improvements in neurocognition across late childhood and early adolescence using neural (Ordaz et al., 2013; Paige et al., 2024) and behavioral (Ahmed et al., 2022; Paige, Ramer, et al., 2026; Tervo-Clemmens et al., 2023; Weigard, Suzuki, et al., 2024; Wiker et al., 2024) measures. These results likewise corroborate prior findings of individual differences and decreases in early adolescent attention problems (Barch et al., 2021; Wiker et al., 2024). Our models also showed that change depended on the initial level of the measure with a negative correlation between baseline and change in all models. This negative correlation is commonly observed in developmental science and may be because adolescents with low initial scores or neural activation have a greater potential to obtain higher scores/activation at subsequent assessments (Rogosa et al., 1982; Von Soest & Hagtvet, 2011).

Regarding our first hypothesis, we found strong support for the co-development of task-control network activation and cognitive efficiency, measured as the drift rate parameter of DDMs fit to individuals’ n-back performance, across ages 10 to 12. Specifically, high task-control network activation was related to faster drift rate at age 10. This is consistent with our previous cross-sectional work linking task-control network activation to better cognitive performance (Paige, Molloy, et al., 2026; Weigard, Angstadt, et al., 2024). We extend this work by demonstrating that development of these brain and behavioral processes support each other across late childhood and early adolescence. Indeed, findings revealed that high task-control network activation at age 10 predicted improvement in drift rate across ages 10 to 12. A reciprocal effect was also observed: faster drift rate at age 10 predicted increases in task-control network activation across time. Moreover, the change scores were associated, such that improvement in drift rate were related to increases in task-control network activation across ages 10 to 12. The associations between baseline levels of task-control network activation and cognitive efficiency were large, consistent with prior work (Paige, Molloy, et al., 2026; Weigard, Angstadt, et al., 2024). The change score associations were in the moderate range of conventional effect size benchmarks and the relations between baseline levels and later change in each variable were far smaller. This corroborates the idea that both task-control network activation and cognitive efficiency show considerable rank-order stability across development, even as they show developmental increases. Indeed, past work has demonstrated the trait-like nature of both cognitive efficiency (Schubert et al., 2016; Tomlinson et al., 2025; Weigard, Suzuki, et al., 2024) and task-control network activation (Paige, Molloy, et al., 2026). Yet, when changes in the rank ordering in each construct do occur, the current results suggest that baseline levels of the other construct play a clear role. In summary, our findings indicate that both neural activation in task-control frontoparietal and dorsal attention networks and cognitive efficiency increase across late childhood to early adolescence, and they do so in parallel.

Regarding the co-development of task-control network activation and attention problems, results provided some support for our first hypothesis. Fewer attention problems at age 10 were concurrently related to high task-control network activation, and they also predicted improvement in task-control network activation across ages 10 to 12. This is consistent with prior work linking high neural activation in the task-control network to fewer attention problems (Paige, Molloy, et al., 2026). However, task-control activation at baseline did not show a statistically significant association with decreases in attention problems across time. The change scores were not significantly associated, suggesting that maturation in task-control network activation did not co-develop with teacher-reported attention problems.

Finally, findings provided strong evidence for co-development between cognitive efficiency and attention problems, in support of our second hypothesis. Indeed, better cognitive efficiency was concurrently related to fewer attention problems at age 10; fewer attention problems at age 10 predicted increases in cognitive efficiency across time; and the change scores were related in the expected direction, such that gains in cognitive efficiency were related to decreases in attention problems. These results corroborate prior work demonstrating that cognitive efficiency estimates from a different ABCD task (stop-signal) showed co-development with parent-reported ADHD symptoms (Wiker et al., 2024). We extend this work by interrogating the neural underpinnings of these associations and highlight that, although the direct links between developmental change in task-control network functioning and attention problems were not strongly supported, developmental changes in cognitive efficiency may mediate associations between task-control network development and attention problems. Future work using three or more repeated assessments should further explore this possibility.

Taken together, these longitudinal findings extend multiple theoretical frameworks of cognitive neurodevelopment. Leading models have traditionally focused on how developmental changes in underlying neural substrates support cognitive maturation across adolescence (Blakemore & Choudhury, 2006; B. Casey et al., 2005; Luna et al., 2015). In addition to providing support for this hypothesis, our study highlights a possible reverse pathway: baseline cognition is associated with subsequent brain development. One possibility is that youth who encode goal-relevant information more efficiently may engage differently with their learning opportunities and broader environments in ways that influence brain development. Another possibility is that higher cognitive efficiency at baseline is a potential marker of earlier neurocognitive maturation, shaping the magnitude of subsequent neurodevelopmental change. These considerations are consistent with early theory emphasizing reciprocal transactions among multidimensional developmental processes (Sameroff, 2009; Sroufe & Rutter, 1984), while highlighting the utility of extending these models to the field of developmental cognitive neuroscience.

### 4.2. Family income, but not family conflict or perceived neighborhood safety, predicts task-control network maturation

Results provided little support for our third hypothesis that low family conflict and high perceived neighborhood safety would be associated with improved task-control network maturation. We found no significant associations between family conflict and task-control network maturation. This is somewhat inconsistent with previous findings from a study using the ABCD baseline sample, which demonstrated that higher family conflict was associated with altered patterns of functional connectivity within and between the frontoparietal, dorsal attention, default mode, and cingulo-opercular networks (Tsomokos et al., 2025). Further, results diverge from a broad literature documenting associations between negative parenting and altered neurocognitive development (Atherton et al., 2020; Blume et al., 2025; Michael et al., 2024). Differences in measurement may help explain these discrepancies. First, the ABCD family conflict measure captures aspects of the broader family environment, including child-related factors, rather than isolating parenting behaviors specifically, which may be less directly related to offspring neurocognitive development. To our knowledge, this was the first study to examine prospective effects of family-level adversity on a neural measure of goal-directed cognition that has demonstrated good psychometric properties (e.g., longitudinal measurement invariance, stability, and construct validity; (Paige, Molloy, et al., 2026). These properties strengthen the interpretability of observed developmental change in goal-directed cognition and underscore the importance of using psychometrically robust neural measures in future research examining how environmental experiences shape neurocognitive development.

High perceived neighborhood safety at baseline was concurrently associated with high task-control network activation. This is consistent with one prior ABCD study that found associations between neighborhood safety and altered functional connectivity within and across the frontoparietal and dorsal attention networks at baseline (Tsomokos et al., 2025). However, contrary to our hypothesis, we did not find significant associations between perceived neighborhood safety and change in task-control network activation. One possibility is that perceived safety captures a relatively distal, weak influence on neurocognitive development that may operate through more proximal experiences, such as chronic stress or few opportunities for cognitive and social engagement (Evans & Kim, 2013). Relatedly, other dimensions of neighborhood adversity, like exposure to community violence or high levels of air pollution, may be more relevant to neurodevelopment (Butler et al., 2018; Kardan et al., 2026; Mattheiss et al., 2022). Future research incorporating multiple dimensions of neighborhood adversity may help clarify whether and under what conditions neighborhood context is associated with maturation in neural systems supporting goal-directed cognition.

Baseline family income significantly predicted change in task-control network activation across early adolescence. Specifically, youth from the highest income bracket displayed faster increases in task-control network activation across time. This finding supports a growing line of research documenting links between socioeconomic status and prefrontal cortex volumes and frontoparietal activity (*for a review, see* Hyde et al., 2020). Notably, past work has been largely cross-sectional, and we advance this line of research by using a psychometrically sound neural measure to demonstrate that family-level socioeconomic status may be linked to maturation of task-control frontoparietal and dorsal attention networks across time.

### 4.3. Limitations

Findings should be interpreted within the context of several limitations. First, the current study utilized only two waves of data to test study aims. Future studies should use three or more assessments, which will allow for more nuanced modeling of developmental trajectories and approaches that disaggregate between- and within-person effects (Curran et al., 2014). Second, neuroimaging quality-control procedures in ABCD may yield analytic samples that disproportionately represent White and socioeconomically advantaged youth (Cosgrove et al., 2022; Gard et al., 2023). This limits the generalizability of our findings and suggests that environmental adversity exposure may have been restricted. Future studies should evaluate the generalizability of findings on the co-development of this neural signature with cognitive performance and attention problems and examine how environmental factors shape brain development in more racially, ethnically, and socioeconomically diverse samples. Third, future work should explore other aspects of the environment that may be important in shaping the neural underpinnings of adolescent cognitive development, including peer relationships (e.g., friendship quality, victimization experiences) and abuse and maltreatment experiences (Lim et al., 2015; Tsomokos et al., 2025; Wen et al., 2023).

## 5. Conclusions

The current study represents the first empirical investigation of the co-development of task-control network functioning with behavioral outcomes, including cognitive performance and attention problems. Results advance our understanding of the mechanisms underlying improvements in behavioral metrics of cognitive abilities across late childhood and early adolescence that have been reported for decades (Ahmed et al., 2022; Gathercole et al., 2004; Luna et al., 2004; Tervo-Clemmens et al., 2023; Williams et al., 1999). Specifically, our study demonstrates that neural activation in frontoparietal and dorsal attention networks shows quantitative increases across ages 10 to 12, and that these increases are *concurrently and prospectively* linked to better cognitive efficiency. Further, this neural maturation develops in parallel with improvements in cognitive efficiency. We also found some support for co-development of task-control network maturation and decreases in attention problems, although these associations appear to be mediated by the maturation of cognitive efficiency. This longitudinal study was also the first to investigate how environmental experiences may shape the activation in task-control networks across early adolescence using a neural measure that demonstrates longitudinal measurement invariance, stability, and good construct validity (Paige, Molloy, et al., 2026). Findings suggested small effects of family-level socioeconomic factors on task-control network maturation. In summary, our study demonstrates, for the first time, that neural maturation in task-control frontoparietal and dorsal attention networks co-develops with improvements in behavioral cognitive efficiency across early adolescence, and highlights the utility of this neural signature for understanding the neural basis of cognitive control development and the mechanisms through which environmental experiences shape brain and behavior.

