## Supplementary Table 1 for "Longitudinal associations between task-control network functioning, cognition, and the environment in early adolescence"

*Supplementary Table 1*. Results from multivariate latent change score model examining associations between change in task-control network functioning, cognitive efficiency, and parent-reported attention problems.

| Parameter | *β* | *SE* | *p* |
|---|---|---|---|
| TC0 ↔ v0 ($\phi_1$) | 0.53 | 0.01 | <.001 |
| TC0 ↔ att0 ($\phi_2$) | -0.09 | 0.02 | <.001 |
| v0 ↔ att0 ($\phi_3$) | -0.12 | 0.01 | <.001 |
| v0 → ΔTC ($\gamma_1$) | 0.10 | 0.03 | <.001 |
| TC0 → Δv ($\gamma_2$) | 0.08 | 0.02 | 0.002 |
| att0 → ΔTC ($\gamma_3$) | -0.04 | 0.02 | 0.063 |
| TC0 → Δatt ($\gamma_4$) | -0.03 | 0.01 | 0.037 |
| att0 → Δv ($\gamma_5$) | -0.08 | 0.01 | <.001 |
| v0 → Δatt ($\gamma_6$) | -0.02 | 0.01 | 0.029 |
| ΔTC ↔ Δv ($\rho_1$) | 0.33 | 0.02 | <.001 |
| ΔTC ↔ Δatt ($\rho_2$) | -0.03 | 0.02 | 0.264 |
| Δv ↔ Δatt ($\rho_3$) | -0.06 | 0.01 | <.001 |

*Note*. The model provided an acceptable fit to the data ($\chi 2(1106)=13042.21$, $p<.001$, CFI=.903, RMSEA=.030(0.030, 0.031), SRMR=.059). The parameters of interest are provided: ($\phi$) correlation at baseline, ($\gamma$) regression and, ($\rho$) correlation of change scores. 0 denotes baseline assessment. TC=Task-control. v=Drift rate (cognitive efficiency). att=Attention problems.
